# Identification of rare *de novo* epigenetic variations in congenital disorders

**DOI:** 10.1101/250787

**Authors:** Mafalda Barbosa, Ricky S. Joshi, Paras Garg, Alejandro Martin-Trujillo, Nihir Patel, Bharati Jadhav, Corey T. Watson, William Gibson, Kelsey Chetnik, Chloe Tessereau, Hui Mei, Silvia De Rubeis, Jennifer Reichert, Fatima Lopes, Lisenka E.L.M. Vissers, Tjitske Kleefstra, Dorothy E. Grice, Lisa Edelmann, Gabriela Soares, Patricia Maciel, Han G. Brunner, Joseph D. Buxbaum, Bruce D. Gelb, Andrew J. Sharp

## Abstract

Certain human traits such as neurodevelopmental disorders (NDs) and congenital anomalies (CAs) are believed to be primarily genetic in origin. With recent dramatic advances in genomic technologies, genome-wide surveys of cohorts of patients with ND/CAs for point mutations and structural variations have greatly advanced our understanding of their genetic etiologies^1,2^. However, even after whole genome sequencing (WGS), a substantial fraction of such disorders remain unexplained^3^. In contrast, the possibility that constitutive epigenetic variations (epivariations) might underlie such traits has not been well explored. We hypothesized that some cases of ND/CA are caused by aberrations of DNA methylation that lead to a dysregulation of normal genome function. By comparing DNA methylation profiles from 489 individuals with ND/CAs against 1,534 population controls, we identified epivariations as a frequent occurrence in the human genome. *De novo* epivariations were significantly enriched in cases when compared to controls. RNAseq data from population studies showed that epivariations often have an impact on gene expression comparable to loss-of-function mutations. Additionally, we detected and replicated an enrichment of rare sequence mutations overlapping CTCF binding sites close to epivariations. Thus, some epivariations occur secondary to *cis*-linked mutations in regulatory regions, providing a rationale for interpreting non-coding genetic variation. We propose that epivariations likely represent the causative genomic defect in 5-10% of patients with unexplained ND/CAs. This constitutes a yield comparable to CNV microarrays, and as such has significant diagnostic relevance.

---

Epimutations represent a class of mutational event where the epigenetic status of a genomic locus deviates significantly from the normal state, and can be classified into two main types: primary epimutations are thought to represent stochastic errors in the establishment or maintenance of an epigenetic state, while secondary epimutations are downstream events related to an underlying change in the DNA sequence^4^. Both secondary and primary epimutations that originate in the germline will be constitutive events found in all cells. In contrast, primary epimutations that occur post-fertilization may result in somatic mosaicism. Constitutive (*i.e.* non-mosaic) epimutations are known to underlie several genetic disorders that can be identified in blood-derived DNA: 5-15% of patients with hereditary non-polyposis colon cancer present with constitutional *MLH1* promoter methylation^5^, and fragile X syndrome, the most common cause of inherited intellectual disability, results from a secondary epimutation in which hypermethylation of an expanded CGG repeat at the *FMR1* promoter causes transcriptional silencing^6^.

We hypothesized that some cases of ND/CA that remain refractory to conventional sequence-based analysis harbor rare epigenetic aberrations (termed epivariations) that that are associated with dysregulation of normal genome function. We studied a cohort comprising 489 individuals with ND/CA: most had been previously tested by CNV microarray, all had undergone exome sequencing, and some had undergone WGS, yet no putatively pathogenic mutations had been identified. Almost 90% of the patients had a ND, and 65% also had multiple CAs, the majority being congenital heart defects (CHD) (Supplementary Table 1). We hypothesized that this cohort represented an optimal population in which to search for novel pathogenic epivariations since an underlying genomic abnormality was suspected, but many common environmental and genetic causes of ND/CA had been excluded. Methylation profiling in ND/CA samples was performed with the Illumina Infinium HumanMethylation450 BeadChip (450k array). Profiles in each ND/CA sample were compared individually against a control cohort comprising 1,534 unrelated individuals from four publicly available datasets (GSE36064, GSE40279, GSE42861 and GSE53045). We also searched for epivariations in two cohorts of population controls by comparison against this same set of 1,534 individuals: 117 families (GSE56105)^7^ were used to assess the inheritance of epivariations in controls (Supplementary Table 2); 2,711 unrelated individuals (GSE55763)^8^ were used to assess the frequency of epivariations in the general population (Supplementary Table 3). We utilized a sliding window approach to identify epivariations in each sample, defined as 1kb regions containing ≥3 probes showing rare outlier methylation absent in the set of 1,534 common control individuals (see Extended Data Fig. 1 and Online Methods section). After stringent quality control, including removal of loci with clusters of poorly hybridizing probes and extensive manual curation to remove technical and batch effects, we identified a total of 143 epivariations in 114 ND/CA samples (*i.e*., 23% of the probands tested). Twenty percent of the ND/CA cohort carried one epivariation (n=98), while 3% of the individuals tested presented two or more epivariations (n=16) (Supplementary Table 4 and Extended Data Fig. 2). Using PCR/bisulfite sequencing, we performed orthogonal confirmation for 70 epivariations (Supplementary Table 5), yielding a 95% true positive rate. Allelic analysis demonstrated that these epivariations represent large methylation changes specifically on one allele, with most showing two clusters of largely methylated and unmethylated reads occurring in approximately equal proportions (Fig. 1).

**Figure 1.**
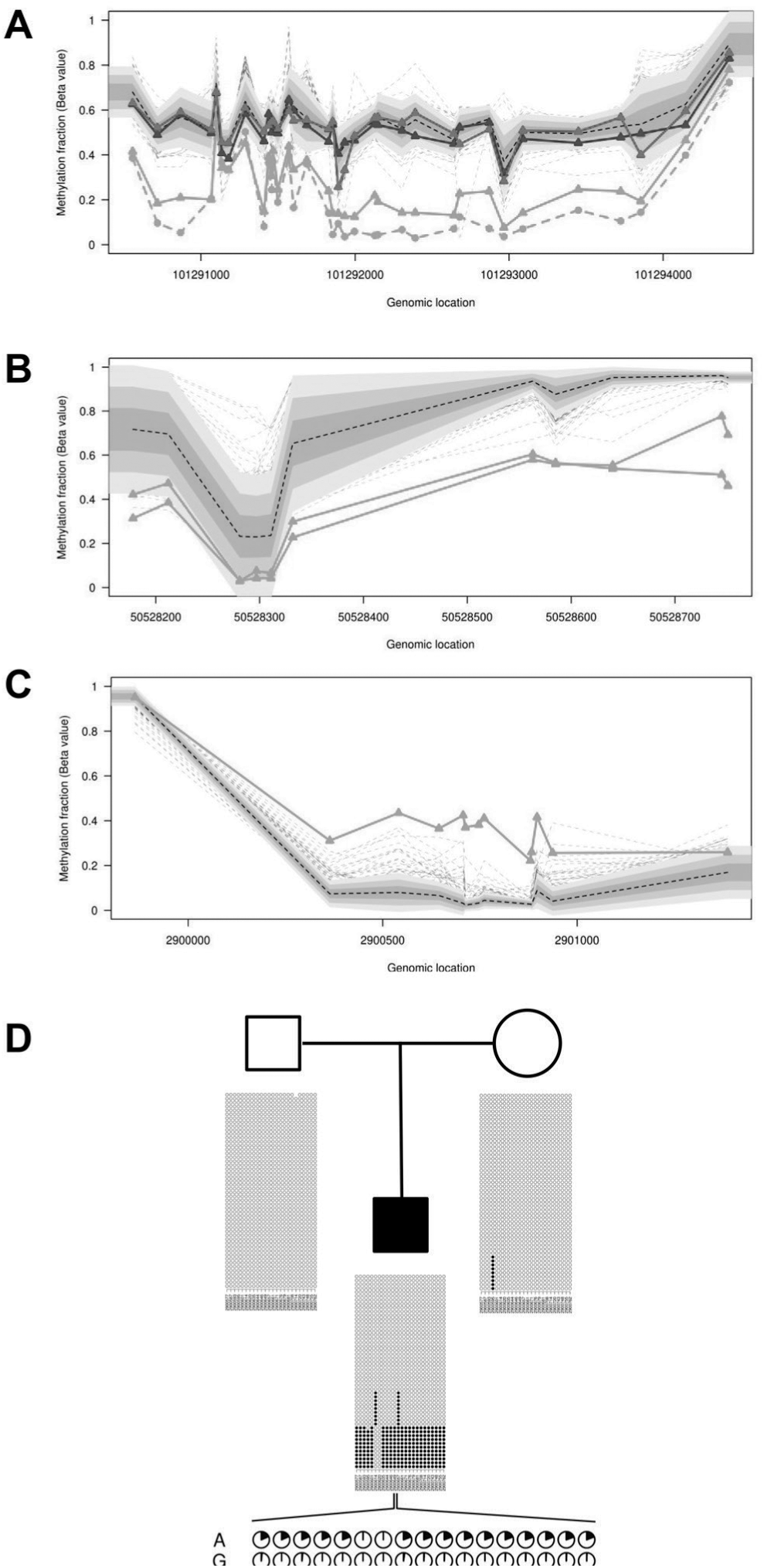
Large gains and losses of DNA methylation identified in patients with ND/CA. Plots A, B and C show β values obtained from Illumina 450k array for probands (highlighted in green) and 1,534 controls (shades of grey corresponding to ±1, ±1.5 and ±2 standard deviations from the population mean, represented by the dashed black line, dashed grey lines represent controls with outlier methylation levels). **A)** Recurrent hypomethylation of the imprinted locus of *MEG3* (hg19: chr14:101290194-101294429) in Proband 398 (solid green line) and Proband 146 (dashed green line). The epivariation in Proband 398 is *de novo* as both mother (red line) and father (blue line) present methylation profiles similar to controls. **B)** Recurrent hypomethylation at the promoter/5’ UTR/first exon of *MOV10L1* (hg19: chr22:50528178-50528751) observed in two unrelated probands: Proband 22 (*de novo* epivariation) and Proband 117 (inheritance unknown). **C)** Hypermethylation of *ZNF57* in Proband 381. **D)** Pedigree and graphical representation of the methyl-seq data consistent with allele-specific nature of a *de novo* hypermethylation identified in *ZNF57* is shown. Each plot shows the methylation pattern for an amplicon, with each row representing a single bisulfite read and each column one CpG in the amplicon. Black circles are methylated CpGs and white circles unmethylated CpGs. Based on the presence of a heterozygous SNP within the DMR (hg19: chr19:2900643), the observed gain of methylation occurs specifically on one allele: each pie chart shows the methylated fraction of reads per CpG.

Epivariations were identified in population controls, and some also occurred in apparently unaffected parents. Thirty three of the 57 DMRs identified in probands for which we investigated inheritance were also present in one parent, and we identified a total of 719 DMRs in the 3,326 control samples analyzed (Supplementary Tables 2 and 3). Twenty four of the epivarations identified in our cases were also present in one or more of these controls, suggesting either that these DMRs are unrelated to the patient phenotype, or perhaps are associated with incomplete penetrance. However, we observed a 1.2 fold enrichment in the frequency of epivariations in the 489 ND/CA samples when compared to 2,711 population controls (Extended Data Fig. 2), although this does not reach statistical significance (p=0.058, two-sided Fisher’s exact test). Testing of parental samples of 57 ND/CA probands showed that 42% (n=24) of the epivariations were *de novo* events. When compared to epivariations found in 117 control pedigrees^7^ (Supplementary Table 2), this represents a 2.8-fold enrichment in the rate of *de novo* epivariations in cases compared to controls (p=0.007, two-sided Fisher’s exact test) (Fig. 2). Thus, while the pathogenic significance of many epivariations is unclear, the paradigm of *de novo* mutational events echoes that observed for other classes of genetic mutation (copy number and single nucleotide variation) deemed pathogenic in ND and CHD cohorts^9,10^.

**Figure 2.**
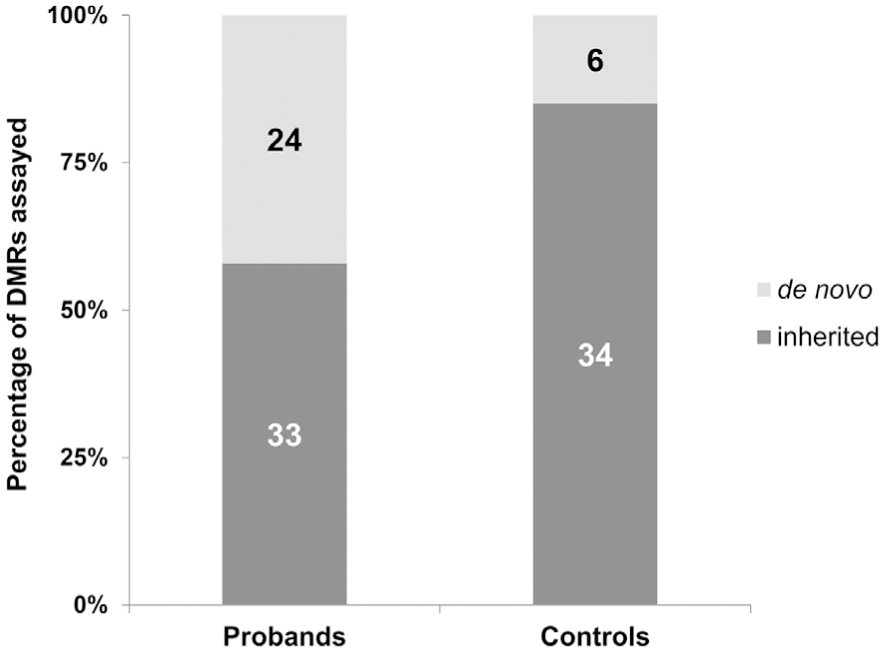
A significant excess of *de novo* epivariations found in patients with ND/CA. We observed a 2.8 fold enrichment for *de novo* epivariations in cases (n=24 out of 57) when compared to controls (n6 out of 40) (p=0.007, two-sided Fisher’s exact test).

In addition to their *de novo* nature, recurrence of mutations found in unrelated patients with a similar phenotype is commonly used as a way of assigning significant evidence for the involvement of a specific gene or locus in disease. We identified 12 recurrent epivariations (Extended Data Fig. 3), *i.e*., the same methylation change was identified in multiple unrelated probands. Of these, two epivariations encompassed the promoters of genes with known disease associations (*MEG3* and *FMR1*)^6,11^, showing our approach successfully detects pathogenic epivariations. A third recurrent epivariation coincides with a locus containing a hypermethylated triplet repeat expansion (*FRA10AC1*)^12^ although this, and four other recurrent epivariations detected in our disease cohort, were also identified in population controls, suggesting that they are unlikely to be pathogenic. One of the novel recurrent epivariations detected only in our patient cohort was found in two patients with CHD (Probands 22 and 117), and represents a recurrent hypo-methylation defect at the promoter/5’ UTR/first exon of *MOV10L1*, a gene with an embryonic heart-specific isoform that interacts with the master cardiac transcription factor NKX2.5^13^ (Fig. 1). Finally, using less stringent criteria for identifying DMRs (see Online Methods), we detected methylation defects in 11 probands at 10 imprinted loci^14^ (Extended Data Fig. 4 and Supplementary Table 6), 90% of which occurred *de novo*. Of note, we observed loss of methylation at two known imprinted loci that have no prior disease associations(*NAA60*/*ZNF597* in Probands 6 and 62, and *L3MBTL1* in Proband 308), although in both cases similar losses of methylation were also observed in population controls, making the pathogenic significance of loss of imprinting at these loci unclear.

Based on previous studies^15,16^, we hypothesized that some epivariations might occur secondarily to an underlying regulatory sequence mutation. In order to identify mutations disrupting regulatory elements (*e.g*., transcription factor binding sites) that might underlie the methylation changes observed in our cohort, we performed high-resolution array CGH and targeted DNA sequencing of 50 DMRs and their flanking sequences. We detected rare sequence mutations that co-segregated with epivariations and potentially impact regulatory elements at 24% of the loci tested: six copy number variations (CNVs) (Fig. 3) and seven single nucleotide variations (SNVs). Where inheritance data from parental samples were available, we found that all of these rare CNVs and SNVs segregated with the presence of the DMR, suggesting that the epivariations occurred secondarily to the underlying sequence mutation.

**Figure 3.**
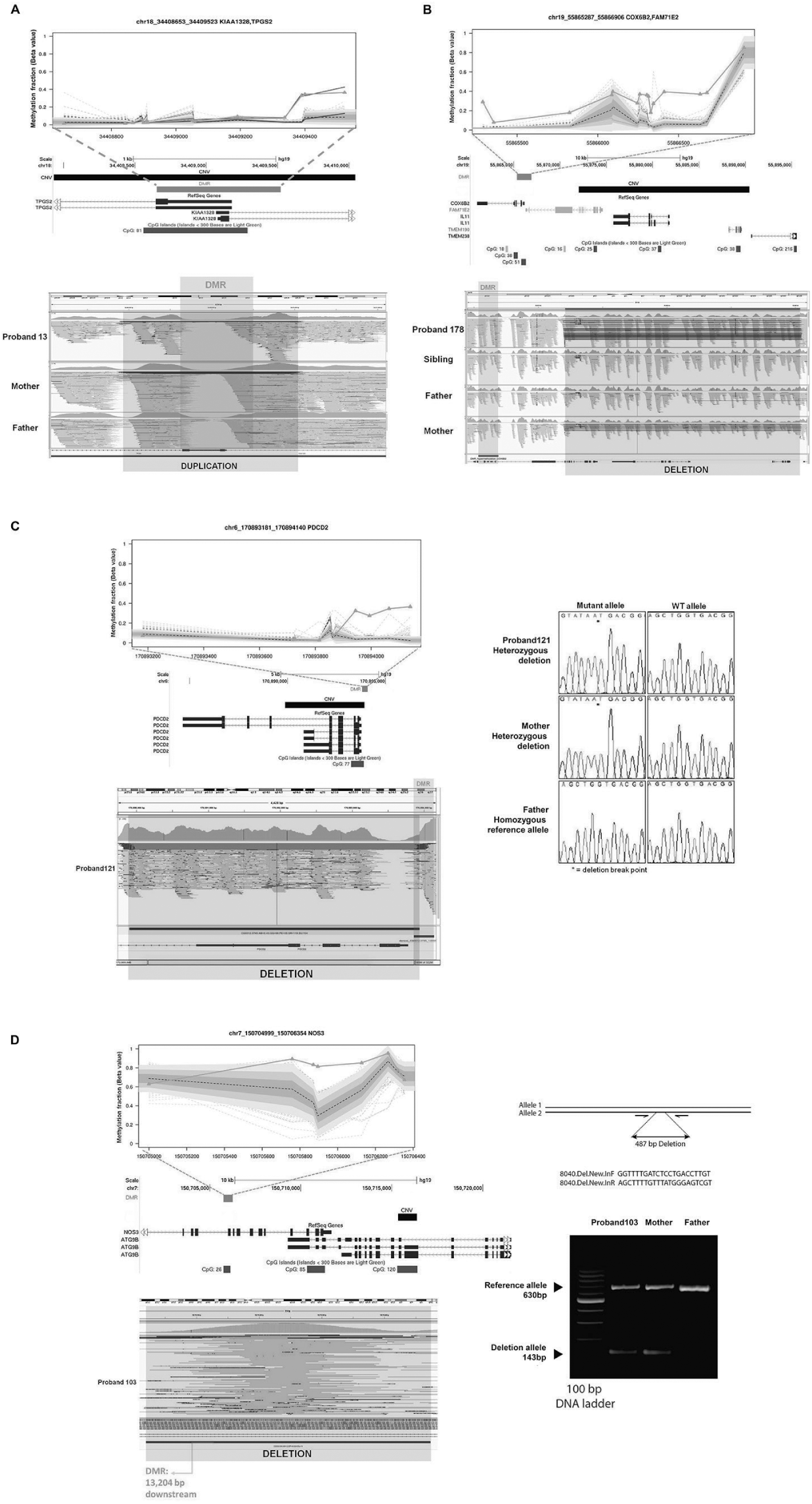
Detection of rare CNVs by targeted sequencing of epivariations and their flanks. Proband 13 carries amaternally inherited DMR at the *KIAA1328/TPGS2* locus. We identified a maternally inherited 2,116 bp duplication that encompasses the DMR. **B)** Proband 178 carries a maternally inherited DMR at the *COX6B2* locus. We identified maternally inherited heterozygous 18,387 bp deletion downstream of the DMR. **C)** Proband 121 carries a maternally inherited DMR at the *PDCD2* locus. We identified a maternally inherited heterozygous 4,061bp deletion flanking the DMR. **D)** Proband 103 carries a maternally inherited DMR at the *NOS3* locus. We identified a maternally inherited heterozygous 487bp deletion located 13,204bp upstream of the DMR.

Of the rare segregating SNVs detected at DMRs, three were SNVs within the canonical binding sites for CTCF (CCCTC-binding factor), a transcription factor with roles in chromatin organization (Fig. 4), including a *de novo* SNV that disrupts a CTCF binding motif in association with a *de novo* epivariation (Proband 70) (Extended Data Fig. 5). In each case, the disrupted CTCF motif was either overlapping, or very close to (separation <1kb), the DMR. This represents a significant enrichment for rare SNVs disrupting CTCF binding sites in the vicinity of epivariations when compared to the same regions in other sequenced samples who did not carry epivariations at these loci (p=0.0015, two-sided Fisher’s exact test), strongly implicating rare *cis*-linked variants in regulatory sequence as a causative factor underlying some epivariations. Furthermore, given the low frequency of *de novo* SNVs and epivariations in the genome, it is highly unlikely that a *de novo* SNV and a *denovo* epivariation would co-occur at the same locus in an individual by chance, providing additional support that some epivariations represent secondary events caused by disruption of CTCF binding. Using paired methylation and sequence data from 90 individuals studied by the 1000 Genomes project (Supplementary Table 7), we replicated this enrichment for rare SNVs disrupting CTCF binding motifs around epivariations (p=0.049, two-sided Fisher’s exact test), identifying two rare CTCF-disrupting SNVs, one of which co-segregates with the presence of an epivariation in multiple unrelated individuals. Though readily detectable by WGS, there is considerable difficulty in interpreting the functional significance of variants outside of coding regions. Thus, we propose that the use of epigenome profiling represents a complementary approach that can provide a rationale for interpreting non-coding genetic variation.

**Figure 4.**
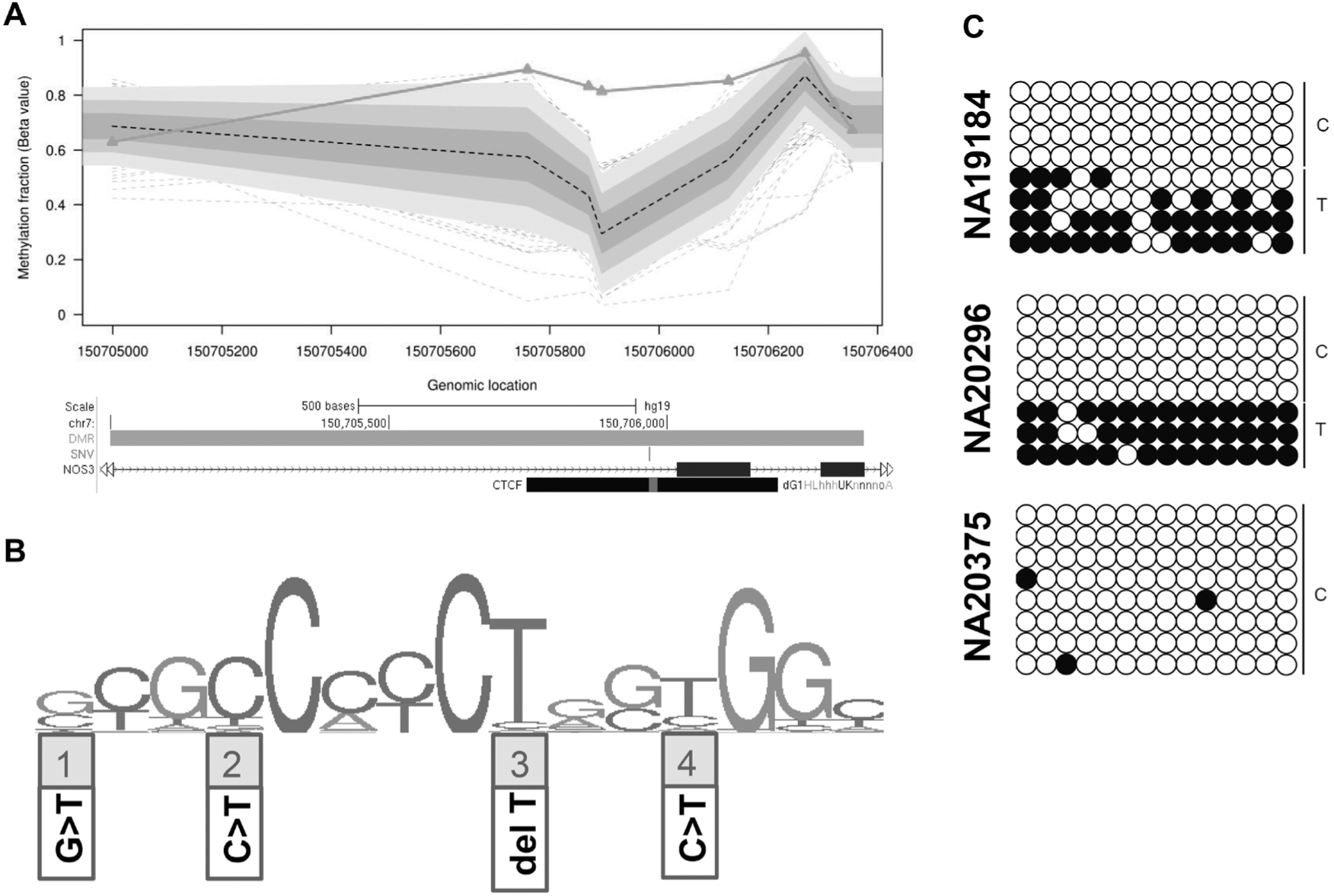
Targeted sequencing of epivariation loci identifies a significant enrichment of rare SNVs within the CTCF canonical binding motif. (A) Hypermethylation in *NOS3* (chr7:150704999-150706354) in Proband 103 (outlier in green); In the lower UCSC Genome Browser view, the DMR location is shown as a green bar, and a rare SNV that lies within the CTCF binding motif (blue region within black bar) in this same individual is shown in red. (B) CTCF motif according to ENCODE Factorbook repository. Rare SNVs overlapping this CTCF binding motif were identified in four DMR carriers: 1) Proband 103: SNV (chr7:150705968 G>T), 2) Proband 70: SNV (chr19:295321 C>T), 3) Proband 176: 1bp deletion (chr20:36793857 delT), 4) HapMap samples NA19239, NA19184; NA20296: rs116767319 (chr5:177707147 C>T). (C) 450k array analysis identified a DMR in NA19239, and a rare SNV (rs116767319) within a CTCF binding motif *in cis*. We tested two other carriers of rs116767319 (NA19184 and NA20296) using allele-specific bisulfite sequencing, and found that both showed methylation on the T allele, thus confirming segregation of the epivariation with SNV. In contrast a sample (NA20375) homozygous for the reference C allele is unmethylated.

**Figure 5.**
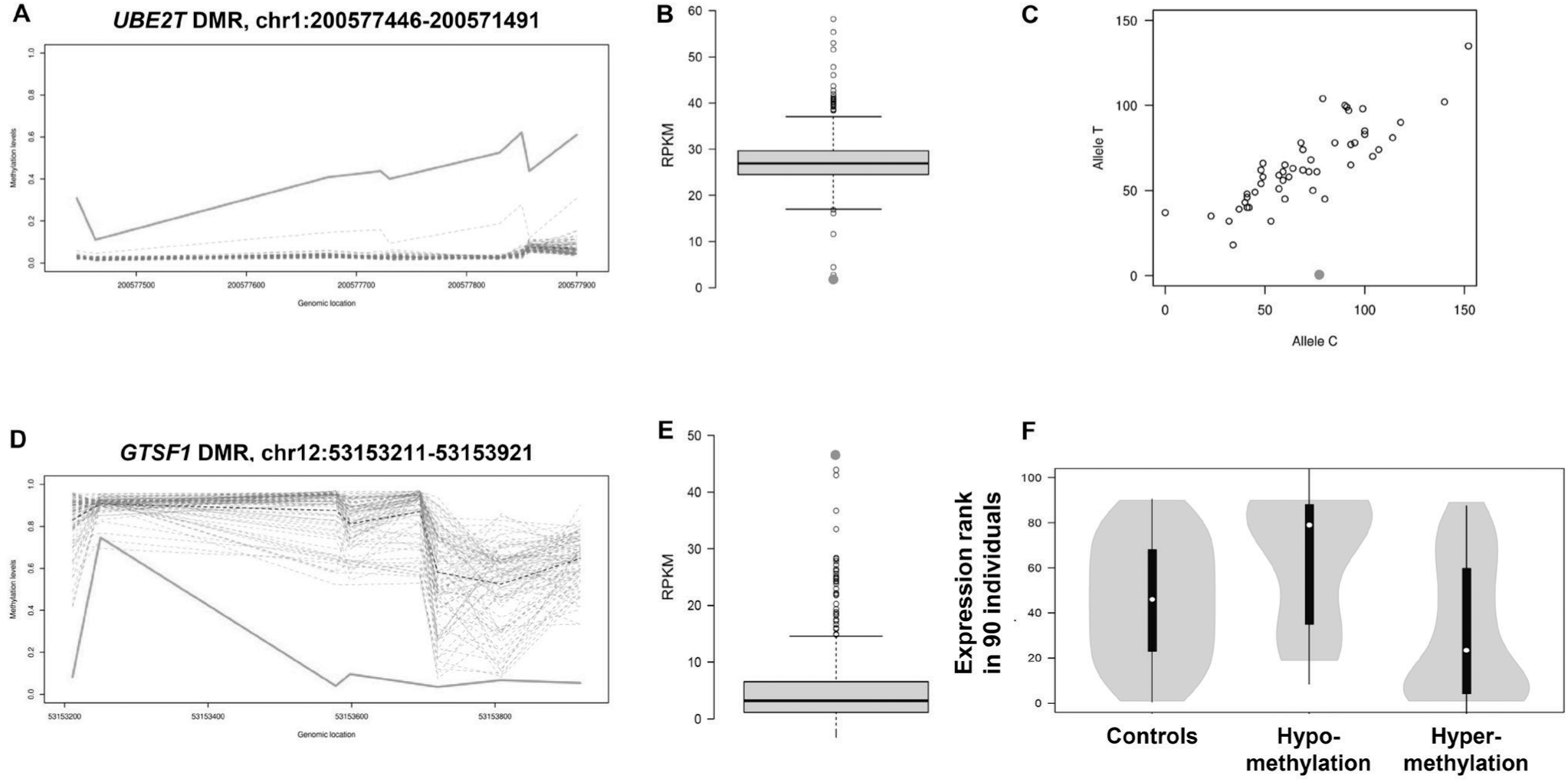
Epivariations are frequently associated with large changes in gene expression. We identified epivariations in 90 lymphoblastoid cell lines studied as part of the 1000 Genomes project, and combined these with SNP genotypes and RNAseq data from a total of 462 samples to measure quantitative and allelic effects of epivariations on gene expression. (**A)** An individual with hypermethylation of the *UBE2T* promoter (solid green line) compared to 89 other individuals (dashed grey lines) presented (**B**) the lowest gene expression (green dot on the boxplot) of the cohort. (**C**) Using heterozygous SNPs within RNAseq reads we observed monoallelic expression of *UBE2T* in the epivariation carrier (outlier highlighted in green). (**D**) An individual with hypomethylation of the *GTSF1* promoter (solid green line) presents (**E**) the highest level of expression (green dot on the boxplot). (**F**) Violin plots show that individuals with hypomethylated epivariations at gene promoters show significantly increased expression of that gene, whereas individuals with hypermethylated promoter epivariations show significantly reduced expression of that gene (p=9.2 × 10^−5^, Wilcoxon Rank-Sum test). In box plots (B and E), the center line shows the median; box limits indicate the 25^th^ and 75^th^ percentiles; whiskers extend 1.5 times the interquartile range from the 25^th^ and 75^th^ percentiles; outliers are shown as individual points. In the violin plot (F), the white dots show the median; box limits indicate the 25^th^ and 75^th^ percentiles; whiskers extend 1.5 times the interquartile range from the 25^th^ and 75^th^ percentiles.

In order to provide insight into the biology and functional consequences of epivariations^17^, we performed studies of gene expression, inheritance and tissue conservation using population datasets of DNA methylation (Supplementary Table 8), gene expression (Supplementary Table 9) and genotype data^18–21^. Using paired RNAseq and DNA methylation data in 90 samples from the 1000 Genomes Project, we verified that epivariations encompassing gene promoters were often associated with large changes in gene expression, with hypomethylation leading to increased expression and hypermethylation to transcriptional repression, consistent with the known repressive effects of promoter DNA methylation (p=9.2 × 10^−5^, Wilcoxon Rank-Sum test) (Fig. 5)^22^. We also observed that many hypermethylated epivariations at promoters are associated with complete silencing of one allele (Extended Data Fig. 6), and, thus, have an impact comparable to that of loss-of-function coding mutations.

**Figure 6.**
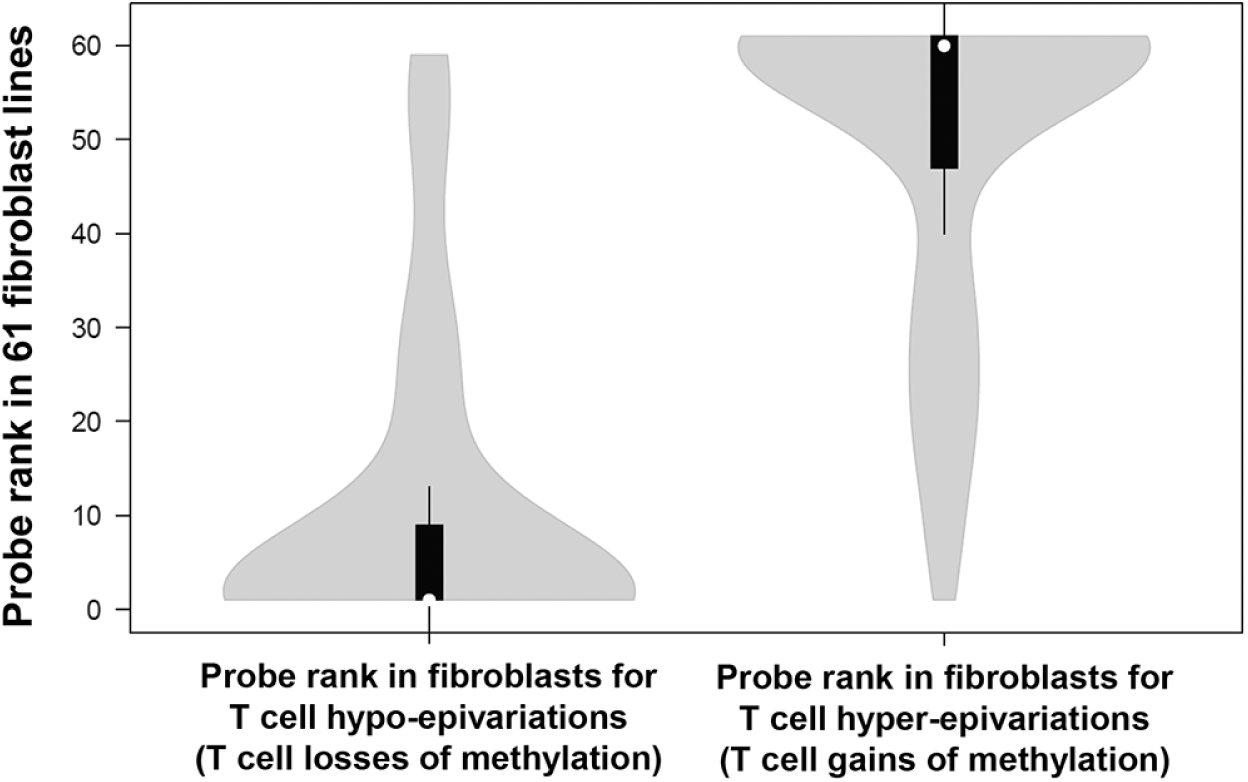
Epivariations detected in blood cells are conserved in fibroblasts from the same individual. The presence of outlier methylation changes in T cells is strongly correlated with outlier methylation in fibroblasts from the same individual (Spearman rank correlation 0.75, p=1.2 × 10^−27^, Wilcoxon Rank-Sum test). White dots show the median; box limits indicate the 25^th^ and 75^th^ percentiles; whiskers extend 1.5 times the interquartile range from the 25^th^ and 75^th^ percentiles.

While epigenetic profiles can vary substantially between cell types^23^, it is unclear whether similar cell-specific variability exists for epivariations. To address that, we analyzed cohorts where methylation profiles were available from multiple different tissues^21^. In samples from the GenCord population, in which methylation data from fibroblasts, B cells and T cells are available, by first identifying DMRs in T cells, we observed a very strong concordance for outlier methylation at the same locus in fibroblasts derived from the same individual (Spearman rank correlation of 0.75, p=1.2 × 10^−27^, Wilcoxon Rank-Sum test) (Fig. 6). Similar concordance for outlier methylation at epivariations was also observed between fibroblasts and B-cells (data not shown).

A similar trend for conservation of epivariations across multiple different post-mortem tissues was also observed in a second cohort^24^. Here, epivariations found in blood were nearly all visible in multiple other somatic tissues sampled from the same individual (Extended Data Fig. 7). Thus, we conclude that the majority of epivariations are constitutive events found in multiple tissues. This provides confidence that epivariations of relevance for ND/CA can be detected using DNA extracted from readily available sources such as peripheral blood leukocytes.

Despite strong evidence that some of the epivariations we observed are secondary events related to the presence of an underlying sequence change (Figs. 3 and 4), we were unable to detect *cis*-linked sequence mutations associated with the majority of epivariations in our cohort, suggesting that these might represent primary epivariations that arose sporadically. As the mammalian genome undergoes several rounds of demethylation and remethylation during gametogenesis, embryonic and somatic development^25^, theoretically there is considerable potential for primary epivariations to be reset to the default state. We therefore assessed how often epivariations are stably transmitted between parents and their offspring. Using a large control cohort comprising 117 nuclear families^7^, we studied the heritability of epivariations between generations, identifying 47 epivariations segregating within these pedigrees. We observed a marked deviation from the expectations of Mendelian inheritance, with only 32 instances of parent-child transmission in 95 informative meioses; significantly fewer than the Mendelian expectation of 47.5 transmissions, (p=0.027, two-sided Fisher’s exact test) (Supplementary Table 2). Therefore, this apparent reduction in heritability indicates that primary epivariations often exhibit non-Mendelian inheritance, and suggests they are frequently reset between generations by epigenetic reprogramming^26,27^.

Our study shows for the first time that epivariations are a relatively common feature in the human genome, that some are associated with changes in local gene expression, and raises the possibility that they may be implicated in the etiology of developmental disorders. In an era when WGS is being applied to many thousands of human genomes, epivariations represent a class of genetic variation that remains undetectable by purely sequence-based approaches. We anticipate that future studies exploring the relationship between sequence variation and epigenetic state will further illuminate the regulatory architecture of the human genome, providing novel insight into the consequences of non-coding mutations.

## Supplementary Information

is available in the online version of the paper.

## Acknowledgements

The authors are grateful to the patients and families who participated in this study and to the collaborators who supported patient recruitment. This work was supported by NIH grant HG006696 and research grant 6-FY13-92 from the March of Dimes to A.J.S., grant HL098123 to B.D.G. and A.J.S., Gulbenkian Programme for Advanced Medical Education and the Portuguese Foundation for Science and Technology (SFRH/BDINT/51549/2011, PIC/IC/83026/2007, PIC/IC/83013/2007, SFRH/BD/90167/2012, Portugal) to P.M., F.L. and M.B., by the Northern Portugal Regional Operational Programme (NORTE 2020), under the Portugal 2020 Partnership Agreement, through the European Regional Development Fund (FEDER) (NORTE-01-0145-FEDER-000013) to P.M., a Beatriu de Pinos Postdoctoral Fellowship to R.S.J. (2011BP-A00515), and a Seaver Foundation fellowship to S.D.R.. The views expressed are those of the authors and do not necessarily reflect those of the National Heart, Lung, and Blood Institute or the National Institutes of Health. Research reported in this paper was supported by the Office of Research Infrastructure of the National Institutes of Health under award number S10OD018522. This work was supported in part through the computational resources and staff expertise provided by Scientific Computing at the Icahn School of Medicine at Mount Sinai.

## Author Contributions

M.B., R.S.J., P.G., H.G.B., J.D.B, B.D.G. and A.J.S. were leading contributors to the design and analysis of this study; M.B., T.K., D.E.G., G.S., P.M., H.G.B, J.D.B, B.D.G contributed with samples of probands and relatives; M.B., D.E.G., S.D.R., J.R., F.L., P.M., L.V., T.K., G.S contributed with patient clinical/genetic information; P.G., N.P., B.J., C.T.W. and K.C. wrote and performed bioinformatic analysis; A.M.T. analyzed and validated methylation profiles of imprinted loci; W.G. performed library preparation and capture for targeted sequencing; C.T. contributed for Agilent custom designed aCGH; H.M. and L.E. processed the Agilent custom designed aCGH; M.B. and A.J.S. wrote the manuscript, all authors commented on it.

## Author Information

The authors declare not having competing financial interests. Correspondence and requests for materials should be addressed to A.J.S.

